# Colibactin-induced damage in bacteria is cell contact independent

**DOI:** 10.1101/2024.06.21.600066

**Authors:** Emily Lowry, Amir Mitchell

**Affiliations:** Department of Systems Biology, University of Massachusetts Chan Medical School, United States

## Abstract

The bacterial toxin colibactin, produced primarily by the B2 phylogroup of *Escherichia coli,* underlies some cases of colorectal cancers. Colibactin crosslinks DNA and induces genotoxic damage in both mammalian and bacterial cells. While the mechanisms facilitating colibactin delivery remain unclear, results from multiple studies supported a delivery model that necessitates cell-cell contact. We directly tested this requirement in bacterial cultures by monitoring the spatiotemporal dynamics of the DNA damage response using a fluorescent transcriptional reporter. We found that in mixed-cell populations, DNA damage saturated within twelve hours and was detectable even in reporter cells separated from colibactin producers by hundreds of microns. Experiments with distinctly separated producer and reporter colonies revealed that the intensity of DNA damage decays similarly with distance regardless of colony contact. Our work reveals that cell contact is inconsequential for colibactin delivery in bacteria and suggests that contact-dependence needs to be reexamined in mammalian cells as well.

**Importance:** Colibactin is a bacteria-produced toxin that binds and damages DNA. It has been widely studied in mammalian cells due to its potential role in tumorigenesis. However, fundamental questions about its impact in bacteria remain underexplored. We used *E. coli* as a model system to study colibactin toxicity in neighboring bacteria and directly tested if cell-cell contact is required for toxicity, as has previously been proposed. We found that colibactin can induce DNA damage in bacteria hundreds of microns away and that the intensity of DNA damage presents similarly regardless of cell-cell contact. Our work further suggests that the requirement for cell-cell contact for colibactin-induced toxicity also needs to be reevaluated in mammalian cells.

## Introduction

Competitive interactions are prevalent within microbial communities, such as those found in the human gut microbiome (1). A common mechanism facilitating these interactions involves the secretion of toxins that target neighboring microbes. One such toxin is colibactin, which is produced by certain bacteria and can bind to the DNA of nearby cells (2–4). Colibactin can induce damage to various bacterial species (5–7) and can also affect host intestinal cells (3, 8). This genotoxin has been linked to numerous human diseases, including inflammatory bowel disease and colorectal cancers (9–12). While colibactin toxicity in host cells has been thoroughly investigated due to its clinical relevance, its ability to damage bacteria remains underexplored. Investigating colibactin toxicity in bacteria could elucidate how colibactin impacts the microbiota while also damaging host cells. Here, we study spatial and temporal dynamics of colibactin toxicity using the *Escherichia coli* model. We discovered that colibactin does not require cell-cell contact for toxicity, as previously suggested.

Colibactin is encoded by a 54kb genomic region known as the *pks* island. This island contains 19 genes needed to synthesize and export the toxin, including non-ribosomal peptide synthetases and polyketide synthases (4). The island also encodes a cyclopropane hydrolase (*clbS*) that protects colibactin producers from self-inflicted damage (4, 13, 14). The toxin itself contains two cyclopropane rings that alkylate DNA and cause interstrand crosslinks (2, 14–18). ClbS cleaves the cyclopropane rings to deactivate colibactin. The *pks* pathogenicity island is most commonly found in *E. coli* strains belonging to the B2 phylogenetic group and is expressed by both pathogenic and commensal strains (4, 19).

Colibactin toxicity has been broadly studied in mammalian cells due to the clinical relevance of the toxin and its association with various human diseases (2, 8, 11, 17, 18, 20–31). Colibactin toxicity has also been observed in bacteria. Colibactin-producing bacteria lacking the ClbS anti- toxin demonstrated auto-toxicity, which became more pronounced in cells with a deletion in the nucleotide excision repair pathway (13, 14). Colibactin has been shown to target other bacterial species, including several *Staphylococcus* species (5–7), several *Vibrio* species, *Clostridium difficile*, and *Enterobacter aerogenes* (7). One recent study suggested that colibactin toxicity in multiple bacteria species is attributed to prophage excision (5). However, prophage-cured *S. aureus* remain susceptible to colibactin, indicating that toxicity mechanisms beyond prophage induction exist (6).

A key gap in knowledge in the field of colibactin-induced damage concerns its delivery route to target cells. Colibactin instability has further confounded the study of this toxin in the extracellular environment (2, 15, 32, 33). The mechanism of colibactin secretion from producer cells remains largely unknown, but some studies in mammalian cells revealed its presence in outer membrane vesicles (34, 35). Multiple studies in both bacteria and mammalian cells claimed cell-cell contact is critical for potent colibactin-induced toxicity (4–7, 17, 25). Here, we directly tested this requirement by monitoring the spatiotemporal dynamics of DNA damage response in recipient bacteria cells. Using live-cell fluorescent reporters of the DNA damage response, we show that colibactin-induced damage occurs within a few hours of exposure. Using two different experimental co-culturing setups, and both engineered and naturally colibactin producing strains, we discover that colibactin-induced DNA damage is detectable hundreds of microns away from the producing bacteria. Our work reveals that cell contact is inconsequential for colibactin delivery in bacteria and suggests that contact dependence needs to be reexamined in mammalian cells as well.

## Results

### Colibactin induces DNA damage in *E. coli*

To evaluate DNA damage in bacteria neighboring colibactin producers, we constructed a transcriptional reporter in an *E. coli* lab strain (Figure 1A). We cloned into a low-copy plasmid a YFP fused to the *recA* gene promoter, known to respond to DNA damage (36). Response of our reporter to DNA damage was validated with mitomycin-C, a known DNA damaging agent that primarily causes interstrand crosslinks (37) (Supplementary Figure 1). We cloned into the same plasmid a CFP expressed from an EM7 constitutive promoter. The colibactin-producing strain was obtained by transforming the same lab strain with a bacterial artificial chromosome (BAC) expressing the pks pathogenicity island (5) (Figure 1A). The same strain, carrying an empty BAC, was used as control. We refer to these strains as pks^+^ and pks^-^ hereafter. Both strains were transformed with a high-copy plasmid expressing an mCherry fluorescent protein from a constitutive promoter (UV5). Expression of CFP and mCherry from constitutive promoters allowed us to identify producers and target cells within a co-culture population, while expression of the YFP allowed us to quantify the level of DNA damage in target cells.

**Figure 1.**
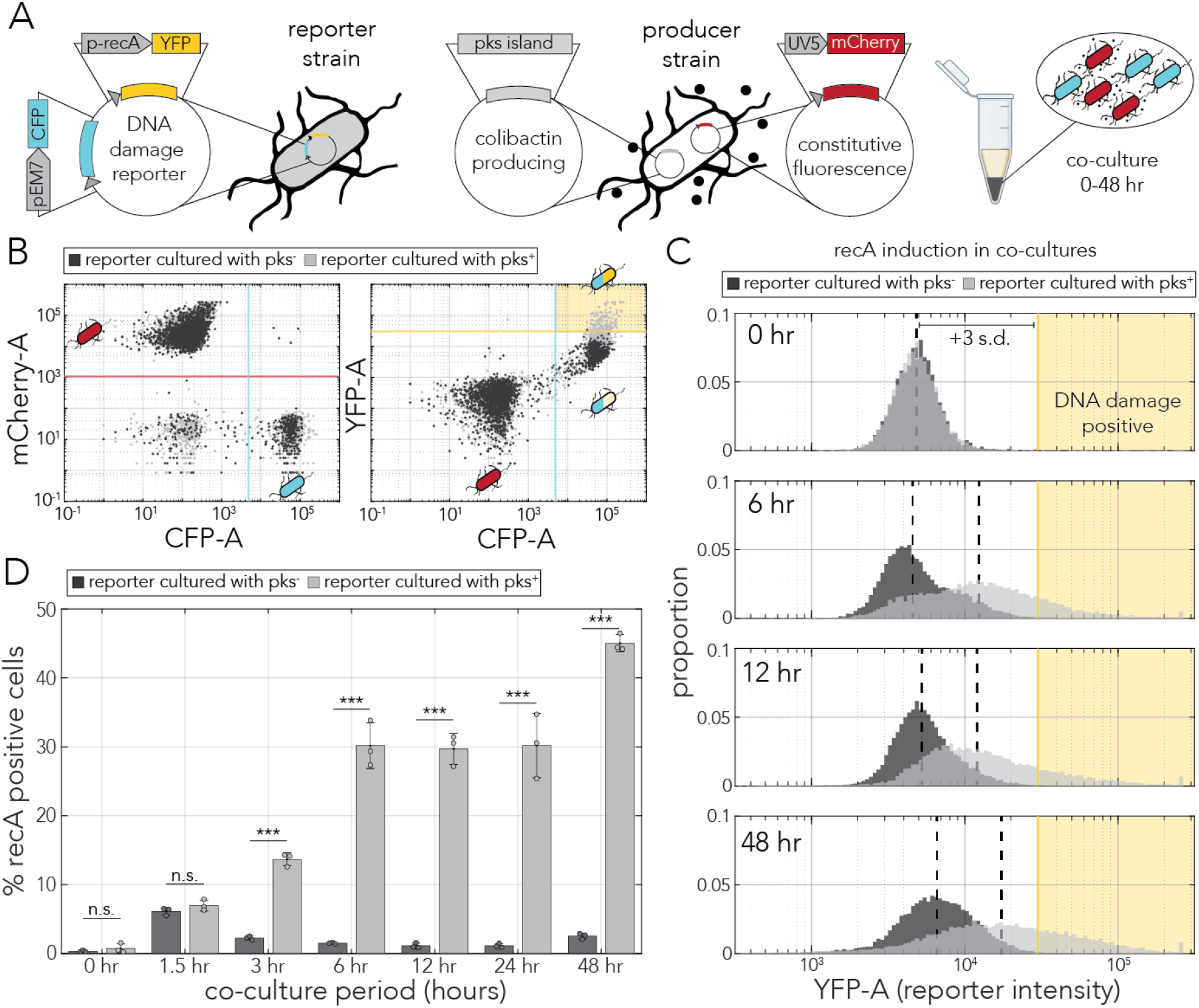
Temporal dynamics of colibactin-induced DNA damage. (A) Plasmid design for tagging reporter cells and monitoring DNA damage response (left) and for tagging colibactin producers (right). Cells expressing each plasmid were co-cultured in a pellet for various time periods. (B) Flow cytometry gating strategy shown with representative cell populations (co-culture for 24 hr). A co-culture of reporters and producers (pks^+^) is marked in light gray and a control co-culture (reporters cultured with non-producers, pks^-^) is marked in dark gray. The left panel shows classification of cells as either reporters (high CFP) or producers (high mCherry). The right panel shows the YFP intensity, and the cutoffs used to identify the proportion of DNA damage positive reporters (yellow shaded rectangle corresponding to high CFP and high YFP signals). (C) Histograms of YFP intensity in reporter cells co-cultured with pks^+^ (light gray) and with pks^-^ (dark gray) over time. The yellow shaded area marks cells positive for a DNA damage response (three standard deviations above the mean signal at t_0_). (D) Percent of DNA damage positive reporter cells when co-cultured with pks^+^ cells (light gray) and pks^-^ cells (dark gray). The error bars mark the standard deviations in three biological replicates and points represent the mean percent positive of each independent replicate (***p<0.001).

We wanted to first test our reporter system and establish the timeline for DNA damage from colibactin in a co-culture model that maximized toxicity prior to testing the requirement for contact. We co-cultured colibactin-producing cells with reporter cells in a pellet at a 10:1 ratio for 48 hours (Figure 1A). We then used flow cytometry to identify populations of producer and reporter cells (Figure 1B, left panel). Using the separated populations, we then quantified the magnitude of the DNA damage response exclusively in reporter cells (Figure 1B, right panel). We chose a conservative threshold, three standard deviations above the YFP baseline, to determine the percentage of DNA damage positive cells (Figure 1B, yellow line).

Using the same threshold for all time points, we were able to evaluate the level of DNA damage in reporter cells over time. Figure 1C shows histograms of reporter fluorescence at selected time points. We observed a clear shift in the intensity histogram in pks^+^ co-cultures, in contrast to almost no shift in pks^-^ co-cultures. Figure 1D shows the percentage of DNA damage positive cells as a function of all tested incubation times. Reporter activity increased within three hours of co-culturing and was maintained for at least two days in 30-50% of cells. In contrast, less than 5% of reporter cells were positive when co-cultured with control pks^-^ cells. We noted that both pks^+^ and pks^-^ co-cultures showed increased reporter activity at very early time points, likely due to the mechanical pressure experienced when pelleting the cells. The results of our experiments agree with the time frame of the colibactin-triggered DNA damage response reported in other bacteria (6) and human cells (4, 17, 25, 26, 28).

### Colibactin induces DNA damage in distant cells

Once we established that colibactin induces detectable DNA damage within a few hours, we leveraged our fluorescent reporter to evaluate the spatial dynamics of toxicity. In this setup, we followed our DNA damage fluorescent reporter in a lawn of reporter cells surrounding a colony of colibactin producers (Figure 2A). We used a fluorescent microscope to quantify the range of colibactin influence around the pks^+^ colony. Figure 2B shows representative microscopy images from these experiments over three selected time points. As the images show, over time we detected increased fluorescence in the reporter channel and a seemingly increase in the range of colibactin influence. However, we also detected increases in the CFP and mCherry channels over time, corresponding to an overall increase in cell number on the plates. We therefore had to devise a metric for quantifying the influence range that can account for overall increase in fluorescence.

**Figure 2.**
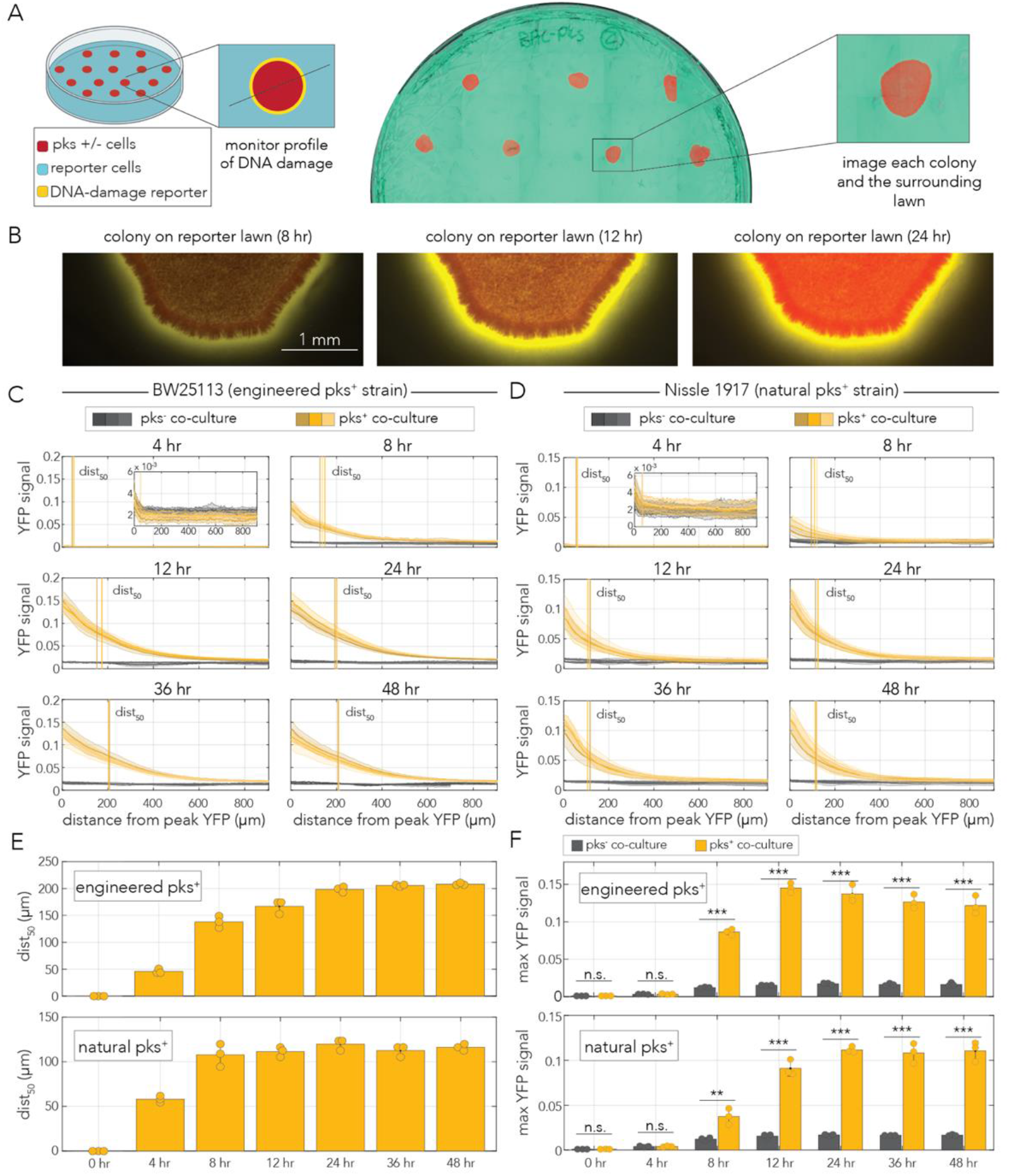
Colibactin induces DNA damage in distant cells. (A) Outline of the experimental setup co-culturing pks^+^ or pks^-^ cells on a lawn of reporter cells. Example image of a reporter lawn expressing CFP with colonies of mCherry-tagged producer cells. (B) Representative microscopy images of a pks^+^ colony tagged with mCherry surrounded by a lawn of reporter cells showing activation of the DNA damage response reporter over time. (C-D) YFP signal decay curves over time in reporter cells co-cultured with the engineered pks^+^ strain (C) or the natural pks^+^ strain (D). The decay curve begins at the peak YFP signal for each technical replicate (N = 10-15). Thick lines represent the mean signal intensity for each biological replicate (n = 3) and shaded areas represent standard deviation. Vertical lines mark the distance from peak YFP signal at which half of the maximal response was observed for each biological replicate (dist_50_). Each biological replicate is colored a different shade of yellow (pks^+^) or gray (pks^-^). (E) Mean distance to half of the maximum YFP signal over time in experiments with engineered pks^+^ cells (top) or with experiments with natural pks^+^ co-cultures (bottom). Error bars represent standard deviation and points represent the mean of each biological replicate (n = 3). (F) Mean maximum YFP signal over time for engineered pks^+^ co-cultures (top) and natural pks^+^ co-cultures (bottom). Control pks^-^ co-cultures are shown in gray and the pks^+^ are shown in yellow. Error bars represent standard deviation and points represent the mean of each biological replicate (n = 3). (**p<0.01, ***p<0.001).

We quantified DNA damage reporter activity along a cross-section that passed from the center of the producer colony (inset in Figure 2A). Profiles of YFP intensity in these cross-sections ranging from the edge of the producer colony and into the lawn of reporter cells reflected the DNA damage signal decay curve. Figure 2C shows the decay curve averaged across 10-15 colonies. The panels represent curves observed at different time points and the individual lines show results from three independent biological replicates. As the panels show, we observed noticeable decay profiles within eight hours of co-culture and an increase in maximal YFP signal in prolonged incubation periods. In the longest incubation periods, elevated YFP signal was clearly noticeable even 500 μm into the reporter lawn. To extract a metric of spatial penetrance that accounts for overall increase in fluorescence, we calculated the distance to 50% of the maximal signal (termed dist_50_). At saturation, the dist_50_ stabilized around 200 μm (vertical lines, Figure 2C). To control for potential artifacts in colibactin delivery due to heterologous expression of the pathogenicity island in engineered cells, we repeated the assay with the Nissle 1917 strain that naturally produces the toxin (Figure 2D). In these experiments we still clearly observed distant DNA damage, although the range of influence was shorter, with dist_50_ stabilizing around 100 μm.

To summarize and compare the spatiotemporal dynamics in the engineered and naturally producing strains, we examined measurements of two key parameters over time. Figure 2E shows dist_50_ in the two strains. In the engineered strain, dist_50_ plateaued after 24 hours and saturated at 208 μm while in the natural producer dist_50_ plateaued already after 8 hours and saturated at 116 μm. The earlier saturation around the natural producer likely arises from lower colibactin expression level in this genetic background. Compatible with this explanation, is the lower signal intensity found at the border of the lawn and the colibactin-producing colony (Figure 2F). At saturation, the maximum YFP signal was higher around the engineered producer by almost 50%.

### Colibactin-induced DNA damage is independent of cell contact

Given our observation that colibactin induces DNA damage hundreds of microns away from the pks^+^ colony (Figure 2C-E), we hypothesized that colibactin may induce DNA damage in neighboring, yet clearly separated, colonies. Such an observation will refute the possibility that DNA damage depends on communication between contacting cells. We therefore designed an additional assay for evaluating DNA damage in both contacting and non-contacting colonies, separated by different distances, on an agar plate. Figure 3A shows an overview of the experimental design: a co-culture of producer and reporter cells were spread on agar plates and allowed to form colonies overnight before visualizing them with a fluorescent microscope. To quantitatively characterize the DNA damage response, we measured the reporter activity profile along a cross-section that passes through the centers of a producer and reporter colony pair (inset in Figure 3A).

**Figure 3.**
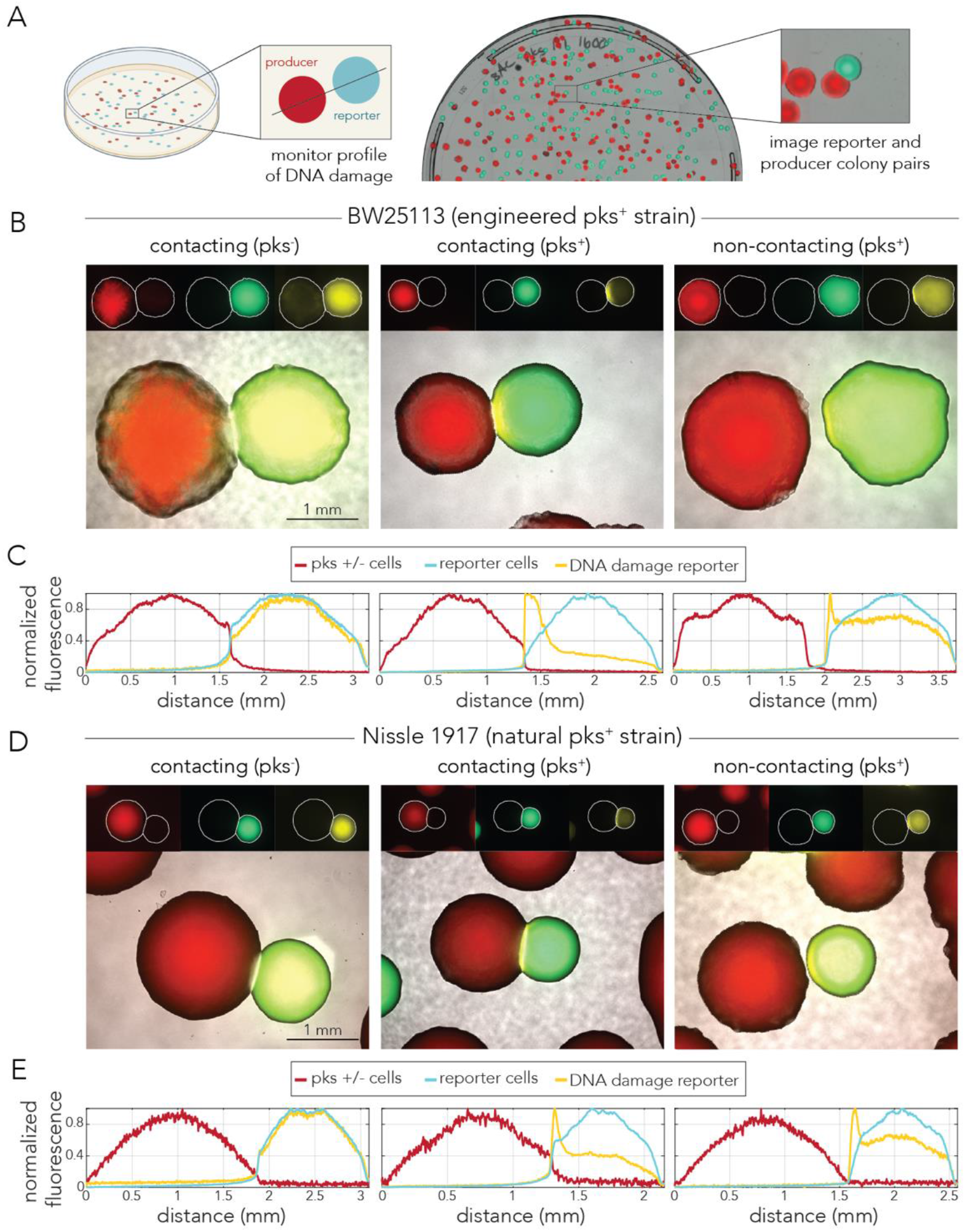
Colibactin-induced DNA damage is cell-contact independent. (A) Outline of the co-culture colony assay. Representative image of a co-culture plate with mixed colonies. (B) Representative microscopy images of reporter colonies (CFP/YFP) next to engineered (BAC) pks^+^ or pks^-^ colonies (mCherry). The signal for each fluorescent tag was min-max scaled in each representative image. In reporter colonies neighboring pks_+_ colonies, YFP signal peaks in the region closest to the colibactin-producing colony. (C) Min-max scaled fluorescent cross-section profiles for representative colonies shown in B. (D) Representative microscopy images of reporter colonies (CFP/YFP) next to natural (Nissle 1917) pks^+^ and pks^-^ colonies (mCherry). The signal for each fluorescent tag was min-max scaled in each representative image. In reporter colonies neighboring pks^+^ colonies, YFP signal peaks in the region closest to the colibactin-producing colony. (E) Min-max scaled fluorescent cross-section profiles for representative colonies shown in D.

Figure 3B shows representative microscopy images from the co-culture experiment with an engineered colibactin producer. As expected, we did not observe any skew in the YFP signal when the reporter colony was next to a non-producing colony (Figure 3B, left panel). Quantification of fluorescence across all channels, shown in the left panel of Figure 3C, supports this observation. The profiles show that the CFP signal, which corresponds to the position and density of the reporter colony, overlaps with the YFP signal. In contrast, reporter colonies contacting pks^+^ cells showed a high intensity YFP signal along a narrow region of contact (Figure 3B, middle panel) and quantification of YFP signal showed a clear positional skew (Figure 3C, middle panel). Lastly, we also observed a weaker, yet highly reproducible, increase in YFP signal in non-contacting colonies that were in proximity (Figure 3B, right panel). This observation was supported by the quantification of the YFP signal (Figure 3C, right panel). To control for potential artifacts in colibactin delivery due to heterologous expression of the pathogenicity island, we repeated the assay with the Nissle 1917 strain. Representative microscopy images and their matching fluorescence profiles are presented in Figures 3D and 3E, respectively. Similarly to the engineered strain, here we also observed a spike in YFP intensity along the edge of both contacting and non-contacting colonies.

We quantified the strength of colibactin-induced DNA damage as a function of the distance from the colibactin-producing colony. Figure 4A shows the YFP signal averaged across dozens of colonies from three biological replicates for co-cultures with the engineered colibactin producing strain. As expected, YFP signal remained at a low and flat baseline in the reporter colonies contacting pks^-^ colonies (Figure 4A, left panel). In contrast, YFP signal spiked in reporter colonies contacting pks^+^ colonies. To quantify the colibactin influence range, we again used the dist_50_ metric. The average dist_50_ was highly reproducible in biological replicates and averaged at 117 μm from the edge of the producer colony (Figure 4C). In non-contacting reporter colonies (separated by at least 10 μm), we also saw clear and reproducible signal decay curves (Figure 4A, right panel). For these colonies, we inferred the distance that matches 50% of the maximum signal observed in contacting colonies. The average dist_50_ was highly reproducible in biological replicates and averaged at 113 μm (Figure 4C). A two tailed t-test rejected the hypothesis that the average dist_50_ is different for contacting and non-contacting colonies (*p*-val = 0.81).

**Figure 4.**
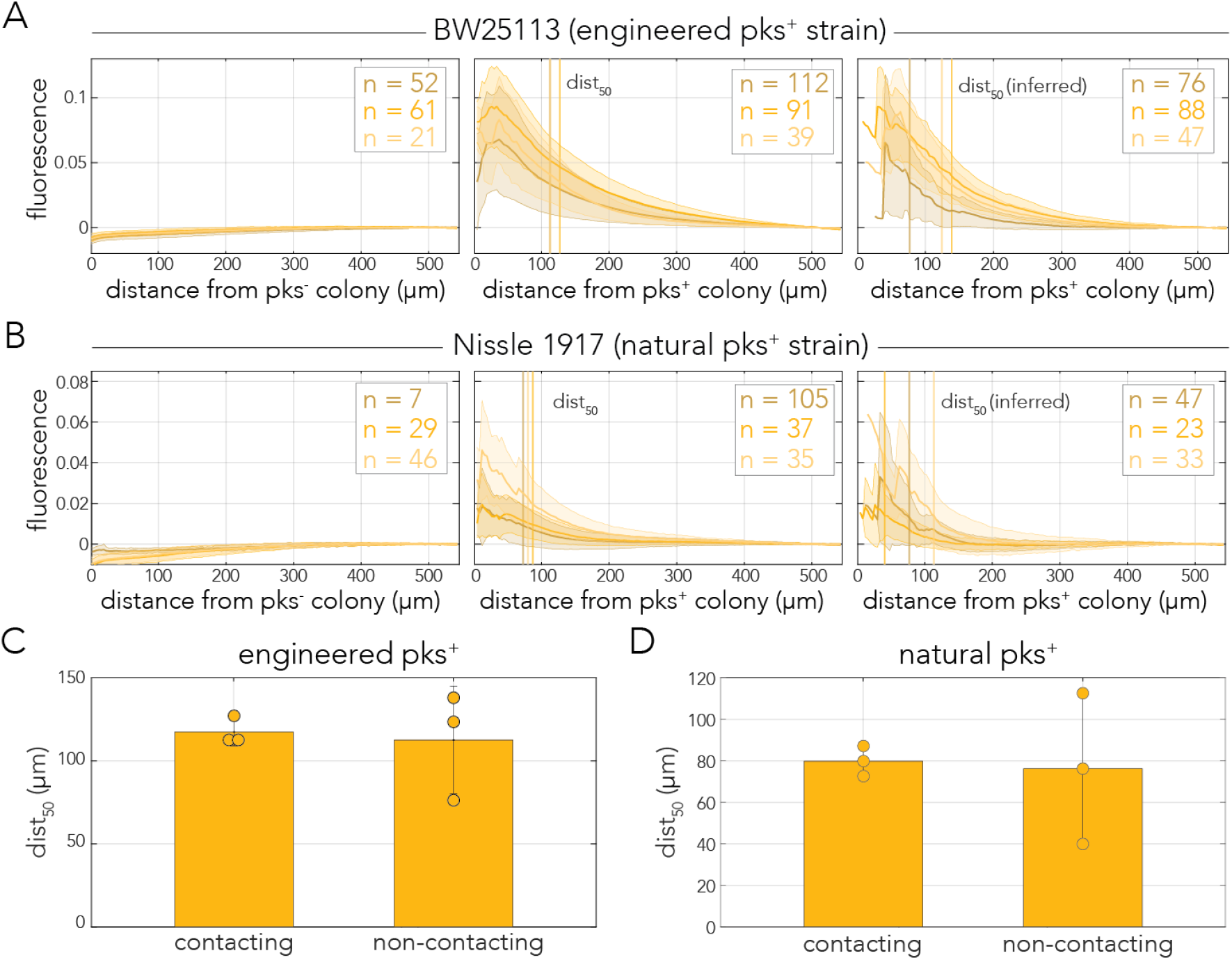
Colibactin-induced DNA damage decay similarly with distance in contacting and non-contacting colonies. (A) YFP decay profiles of reporter colonies plated with engineered cells across three conditions: contacting pks^-^, contacting pks^+^, non-contacting pks^+^. (B) YFP decay profiles of reporter colonies plated with Nissle 1917 cells across the same three conditions. (A, B) YFP signal was aligned to the edge of the mCherry colony and averaged across dozens of colonies in the same plate (n shows the number of colonies monitored). Shaded area marks the standard deviation for each biological replicate. Vertical lines indicate the decay dist_50_ (50% signal) for each biological replicate. (C) Decay dist_50_ calculated and averaged averaged across biological replicates for the engineered strain and (D) for the natural strain dist_50_ in non-contacting colonies is inferred using the averaged YFP intensity measured at dist_50_ for contacting colonies. Error bars represent standard deviation.

We repeated the colony-based assay to examine signal decay in colonies neighboring natural colibactin-producing cells. Figure 4B shows the characteristic decay profiles of reporter colonies neighboring Nissle 1917 cells from three biological replicates. As expected, we observed an overall weaker reporter signal, compared to the engineered strain. The average dist_50_ was highly reproducible in biological replicates of contacting colonies and averaged at 80 μm from the edge of the producer colony (Figure 4D). In non-contacting colonies we observed a higher variation across biological replicates, likely due to the lower level of signal, with an inferred dist_50_ of 76 μm. A two tailed t-test rejected the hypothesis that the average dist_50_ is different for contacting and non-contacting colonies (*p*-val = 0.87).

Lastly, we validated that non-contacting colonies are truly separated from one another and that we are not overlooking pks^+^ bacteria that may have swarmed to the visible reporter colony. We reasoned that if swarming cells were sparse, they may go undetected by low magnification microscopy. We used a fine needle to sample the edge of twelve reporter colonies that showed elevated YFP expression and were separated by 60-300 μm from colibactin-producing colonies. We then resuspended these samples and plated them at different densities on agar plates to check for strain cross contamination (Supplementary Figure 2). We examined plates that had thousands of colonies, with a median of approximately 7500 colonies, for CFP and mCherry signal. In all experiments we exclusively observed CFP-tagged colonies (Supplementary Table 1). We can therefore conclude swarming pks^+^ cells are either absent or extremely rare in the edge of our separated reporter colonies. In summary, our colony-based assay reveals that colibactin-induced damage is evident across clearly non-contacting colonies. Moreover, signal intensity as a function of distance revealed that the signal decay profiles were indistinguishable between contacting and non-contacting colonies.

## Discussion

Colibactin-producing bacteria are not uncommon in the gut microbiome of healthy humans and their increased prevalence is evident in multiple human diseases (9–12). Compelling evidence from colorectal tumors strongly supports the premise that colibactin acts as a tumorigenic mutagen (3, 8). While the clinical relevance of colibactin-induced damage has motivated intense research in mammalian cell cultures, it has left fundamental questions underexplored in bacteria. Addressing these open questions is important for interpreting colibactin’s impact on other members of the host microbiome and potentially also for a deeper understanding of colibactin-host interactions. Here we focused on the spatiotemporal dynamics of colibactin-induced DNA damage in bacteria to directly examine the widely accepted premise of cell- contact dependance (4–7, 17, 25).

We monitored the dynamics of colibactin-induced DNA damage by cloning a transcriptional fluorescent reporter that tracks expression of the *recA* gene, a key factor in the homologous recombination DNA repair pathway (36). By combining this reporter with fluorescent tags that uniquely marked colibactin producers and target cells, we were able to closely track the spatiotemporal dynamics of the DNA damage response in mixed-cell populations. This approach validated previous reports in both bacteria (6) and human cells (4, 17, 25, 26, 28) by showing that DNA damage is already detectable within several hours of colibactin exposure (Figure 1D). However, results from co-culture populations also revealed a seeming discrepancy with previous works by revealing that DNA damage is detectable hundreds of microns away from colibactin-producing cells (Figure 2C-E). The observation of contact independence was reinforced by our colony-based assay that showed that the DNA damage response is triggered in reporter colonies that are clearly separated from colibactin-producing colonies (Figures 3C and 3E). Lastly, quantification of signal intensity as a function of distance revealed that the signal decay profiles were indistinguishable between contacting and non-contacting colonies (Figures 4B and 4E). This observation indicated that cell contact does not alter the amount of DNA damage targeted cells experience beyond what is expected by proximity alone.

Taken together, our results establish that colibactin-induced damage in bacteria is cell-contact independent. Since we observed similar effects, albeit weaker, with a strain that naturally produces colibactin, contact independence is not an inadvertent artifact of the high heterologous expression in genetically engineered cells. Importantly, our conclusion contests the premise that cell-cell contact is required for colibactin toxicity in bacteria (5–7). This seeming contradiction can be rationalized by the different experimental methods that we and others used. Previous works inferred contact dependence from experiments that separated producers and target cells with a small pore membrane in a transwell (5) or by measuring viability in mixed populations growing in a colony compared to a planktonic state (6). Given our conclusion on the effective distance, it seems likely that previous experimental setups were not adequately sensitive for detecting DNA damage at very close proximity. Interestingly, since the conclusion favoring the requirement for cell contact in mammalian cells also relied on a transwell-based assay (4, 17), contact-independent interactions taking place at closer ranges may have gone unnoticed. Our work therefore suggests that the premise of contact-dependence should also be reevaluated in mammalian cells with alternative, and highly sensitive, methods.

## Methods

### Bacteria strains used

| Strain | Nickname | Source | Use |
| --- | --- | --- | --- |
| BW25113 $\Delta gspI::kan$<br>pRecA** | Reporter strain | This study | DNA damage flow cytometry and microscopy |
| BW25113<br>pBeloBAC11+pks-<br>mCherry* | Engineered pks<sup>+</sup><br>mCherry | This study | DNA damage flow cytometry and microscopy |
| BW25113 pBeloBAC11-mCherry* | Engineered pks<sup>-</sup><br>mCherry | This study | DNA damage flow cytometry and microscopy |
| Nissle 1917 (EcN)<br>$\Delta mchDEF::kan$ mCherry | Nissle 1917 pks<sup>+</sup> strain<br>(microcin deletion) | This study | DNA damage microscopy |
| Nissle 1917 (EcN)<br>$\Delta mchDEF::kan \Delta clbN::chl$<br>mCherry | Nissle 1917 pks<sup>-</sup> strain<br>(microcin deletion) | This study | DNA damage microscopy |

\*parental BW25113 pBeloBAC11 and BW25113 pBeloBAC11+pks strains were received from the Balskus lab (5).
\*\*parental BW25113 $\Delta gspI::kan$ strain was from the Keio collection (38).

### Media and growth conditions

All experiments were performed in either Lysogeny Broth (LB) or minimal synthetic media (M9 salts supplemented with 0.4% glucose, 2 mM MgSO_4_, 0.1 mM CaCl_2_, 0.2% amicase). Overnight cultures for all experiments were grown at 37°C with 200 rpm orbital shaking. During the overnight growth of antibiotic-resistant strains, we added antibiotics at the following concentrations: 50 μg/mL spectinomycin, 50 μg/mL kanamycin, 25 μg/mL chloramphenicol, and 50 μg/mL carbenicillin. Agar-based experiments were conducted on M9 1.5% agar.

### Cloning deletion strains

The two microcins MccM and H47 were deleted from Nissle 1917 using lambda red recombination. The insert had 40 bp homology arms to the upstream and downstream genomic regions of the *mchDEF* genes and a kanamycin resistance cassette. *clbN* was deleted from Nissle 1917 using lambda red recombination with an insert containing 40 bp homology arms to the upstream and downstream genomic regions and a chloramphenicol resistance cassette.

### Fluorescent reporter plasmids

We cloned the DNA damage reporter plasmid with the Gibson assembly method (39) using In- Fusion Snap Assembly Master Mix (Takara, cat# 638947). In a single assembly reaction, we integrated a YFP and the *recA* promoter into a plasmid backbone containing a spectinomycin resistance cassette and CFP. When amplifying the backbone, we also replaced the CFP promoter with a 48 bp EM7 promoter that was encoded on the amplification primer. The 81 bp *recA* promoter was amplified from the BW25113 genome (the promoter region was defined according to previous work (40–42)). The final plasmid was a low-copy plasmid with the sc101 origin and spectinomycin resistance. Plasmid assembly was validated with Sanger sequencing spanning the integration sites.

The BAC and Nissle 1917 pks^+^ and pks^-^ strains were cloned to express an mCherry plasmid. The plasmid was high-copy with a colE1 origin and an ampicillin resistance cassette. The mCherry is expressed under the constitutive UV8 promoter.

### Monitoring DNA damage reporter with flow cytometry

Cultures of the reporter strain and engineered pks^+^ and pks^-^ strains containing mCherry expressing plasmids were grown overnight in LB. 1 mL of overnight culture was washed three times in PBS and resuspended in M9 media. Cultures were then diluted 1:200 and allowed to grow for 12 hours. Following growth, optical density at 600 nm (OD_600_) was measured and cultures were diluted to OD_600_=0.1. The reporter culture was then diluted 1:10 into M9 media that contained either the engineered pks^+^ or pks^-^ strain at the same density. The co-culture was maintained in a 96 deep-well plate (Eppendorf, cat# 2231000920) in a final volume of 500 μL. The co-cultures were pelleted at 4000 g for 6 minutes and were then incubated at 37°C without shaking for up to 48 hours. At different pre-determined time intervals, specific wells were resuspended and transferred to a new 96 deep-well plate. Co-cultures were then pelleted at 4000 g for 6 minutes and supernatant aspirated. Cells were fixed with 3.7% formaldehyde for 15 minutes at room temperature. Cells were washed with PBS twice and resuspended in 300 μL PBS. Fixed cells were stored at 4°C for up to three days, then analyzed by flow cytometry. Flow cytometry was performed with BD LSRFortessa with a high throughput sampler. CFP was measured with a 405 nm laser with a 525/50 nm filter and 505 nm long pass filter. YFP was measured with a 488 nm laser with a 530/30 nm filter and 505 nm long pass filter. mCherry fluorescence was measured with a 561 nm filter with a 610/20 nm filter and 600 nm long pass filter.

We analyzed flow cytometry data with a custom MATLAB (MathWorks) script. For gating purposes, unstained, single color, and sample treatment cell populations were overlaid in plots. To separate cell populations from debris, we first gated events using forward scatter and side scatter areas. The filtered events were then further gated to identify single cells with forward scatter area and forward scatter height. Single cells were then gated on each fluorescent channel (by fluorescent channel area). We used a conservative cutoff to classify reporter cells that were positive for the DNA damage response (by YFP signal). We relied on the YFP intensity distribution of the reporter cells at the start of the experiment (t=0 hours) and set the cutoff to be the mean intensity plus three standard deviations.

### Monitoring DNA damage reporter by microscopy in a lawn co-culture

Cultures of the reporter strain, the engineered pks^+^ and pks^-^ strains, and Nissle 1917 pks^+^ and pks^-^ strains (with a microcin deletion, as microcins are also toxic to neighboring cells) were grown overnight in LB with antibiotics. The density of overnight cultures was measured using OD_600_. Strains were then washed twice in PBS and then the reporter strain was diluted to an OD_600_ of 4 and the pks^+^ and pks^-^ strains were concentrated to an OD_600_ of 10. 150 μL of reporter strain was spread on M9 agar plates and allowed to dry. The pks^+^ and pks^-^ strains were spotted in a 1 μL volume and immediately imaged for a 0-hour time point. Plates were incubated at 37°C until each subsequent time point up to 48 hours. Colonies and the surrounding lawn were imaged at 2.5x magnification using CFP (475nm), YFP (524nm), mCherry (610nm), and brightfield channels on a Zeiss Axio Observer.Z1 epifluorescence microscope.

Microscopy image analysis was performed with custom scripts in MATLAB (MathWorks). Briefly, a user marked a line from the center of the producer colony extending into the reporter lawn using the images obtained from the bright-field and CFP channels. The pixel intensity for each fluorescent channel was determined along this cross-section line. Signal decay away from a colibactin producing colony was measured from the peak YFP signal into the reporter lawn. Maximum YFP signal intensity was reported from the peak of the YFP signal. We calculated dist_50_, the distance at which we observed 50% of the maximal response, for contacting colonies by first smoothing each YFP signal decay with a moving window of 20. Decays were smoothed to control for noise in the baseline of the YFP signal, especially at early time points where the overall signal was low. We then found the fluorescent value halfway between the max YFP signal and the baseline signal (YFP_50_). The baseline signal was determined by calculating the average YFP signal of the furthest 20 pixels from the pks^+^ colony. We then found the distance along the decay profile that corresponds to YFP_50_.

### Monitoring DNA damage reporter by microscopy in single colonies

Cultures of the reporter strain, the engineered pks^+^ and pks^-^ strains, and Nissle 1917 pks^+^ and pks^-^ strains (with a microcin deletion, as microcins are also toxic to neighboring cells) were grown overnight in LB with antibiotics. The density of overnight cultures was measured using OD_600_ and were mixed at a ratio of 5:1 or 1:1 (producers to reporters). Mixed co-cultures were then diluted to multiple concentrations and spread on M9 agar plates with glass beads to achieve a range of 400-1,600 colonies per plate. After 24 hours of incubation at 37°C, plates were imaged with a Zeiss Axio Observer.Z1 epifluorescence microscope. Colonies were imaged at 2.5x magnification using CFP (475nm), YFP (524nm), mCherry (610nm), and brightfield channels.

Microscopy image analysis was performed with custom scripts in MATLAB (MathWorks). Briefly, a user marked a line crossing two neighboring colonies and additional lines marking the colony edges on the image obtained from the bright-field and CFP channels. The pixel intensity for each fluorescent channel was determined along this cross-section line. Signal decay away from a colibactin producing colony was measured from the peak YFP signal (at the reporter colony edge) and into the reporter colony itself. For non-contacting colonies, values at positions between the two colonies were ignored. Baseline colony YFP expression was determined by calculating the mean intensity of the furthest 20 pixels from the pks colony in the decay curve. This value was subtracted from each decay curve to control for differences in baseline YFP expression in each colony. The distance between each colony pair was calculated by the user marked colony edges.

We calculated dist_50_ for contacting colonies by first finding the fluorescent value halfway between the max YFP signal and the baseline signal (YFP_50_). We then found the distance along the decay profile that corresponds to YFP_50_. The dist_50_ for non-contacting colonies was inferred by the YFP_50_ value measured earlier for contacting colonies (averaged across all replicates). This was to account for the fact that non-contacting colony decay curves were not peaking at the maximum YFP reporter response.

### Colony cross contamination test

We tested strain purity in the edge of non-contacting colonies to validate that increased YFP intensity is not attributed to swarming pks^+^ cells that were undetectable with the low magnification microscope. This experiment setup was identical to the single colony microscopy assay described above. Non-contacting co-culture colonies were identified on agar plates and the distance separating them was calculated by image analysis as described previously. The colonies were observed under a LEICA S6 E microscope at 2x magnification and the edge of the reporter colony was picked with a flattened platinum wire (forming a fine needle). The collected cells were inoculated into PBS from the wire picker for plating. After picking, we confirmed that only the reporter colony was disturbed at 2.5x magnification on a fluorescent microscope. We excluded any colony pairs where the pks^+^ colony appeared disrupted upon visual inspection at 2.5x on a fluorescent microscope. The suspended reporter colony samples were serially diluted 1:1 for four dilutions (in 1 mL PBS total) before 100 μL from each dilution was plated on LB agar plates. The plates were cultured overnight at 37°C and imaged the following morning with a Canon EOS Rebel T3i camera. Colony counts were calculated with FIJI and CFUs were back-calculated based on the dilution factor. Plates were imaged on a microscope with CFP (475nm) and mCherry (610nm) channels to confirm the presence of only CFP-tagged colonies.

## Data and materials availability

MATLAB codes used in this article are deposited to GitHub (DOI: 10.5281/zenodo.11659009).

## Supplementary Materials

**Supplementary Figure 1.**
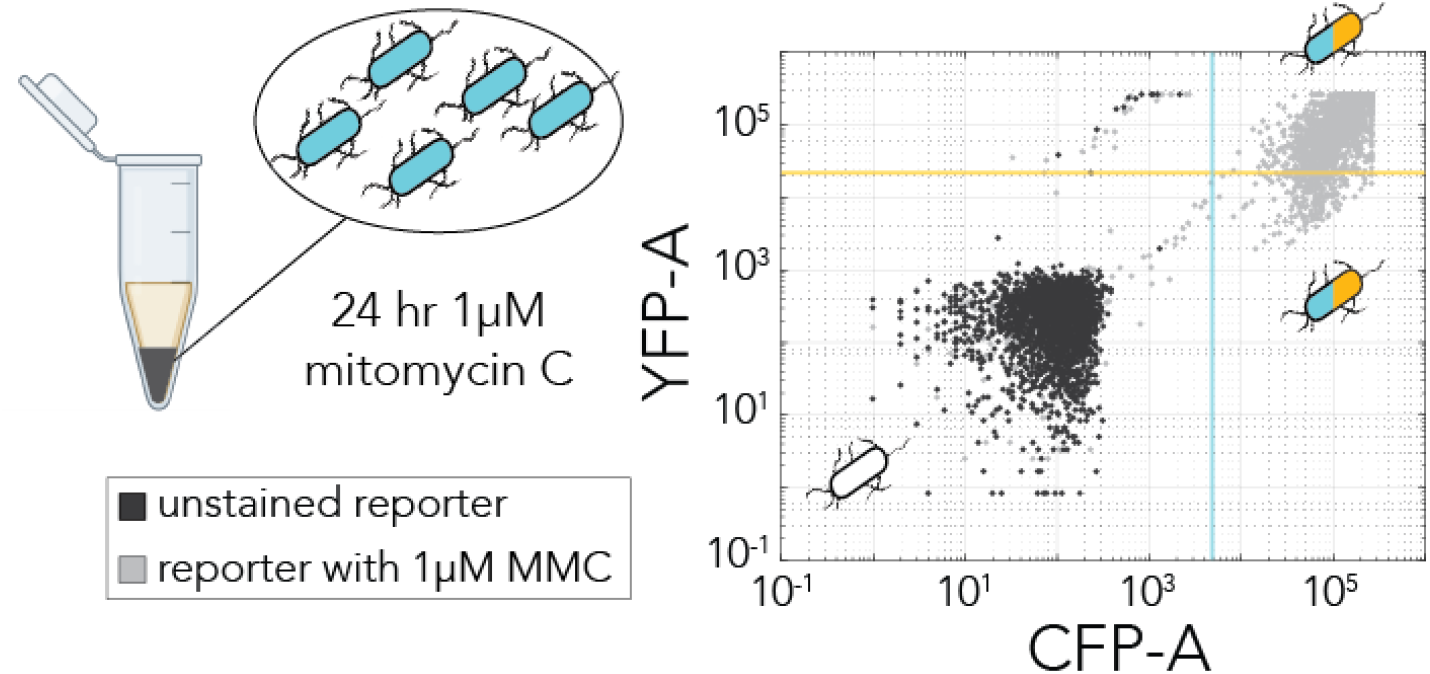
Validation of the recA DNA damage reporter. Bacteria expressing the recA DNA damage reporter plasmid were incubated with 1 μM mitomycin-C (MMC) for 24 hours pelleted in minimal media. *E. coli* cells lacking the fluorescent plasmid served as an unstained control. Both cell populations were analyzed by flow cytometry. The same flow cytometry gating from Figure 1B is displayed on the scatter plot with the unstained population in dark gray and the MMC treated population in light gray.

**Supplementary Figure 2.**
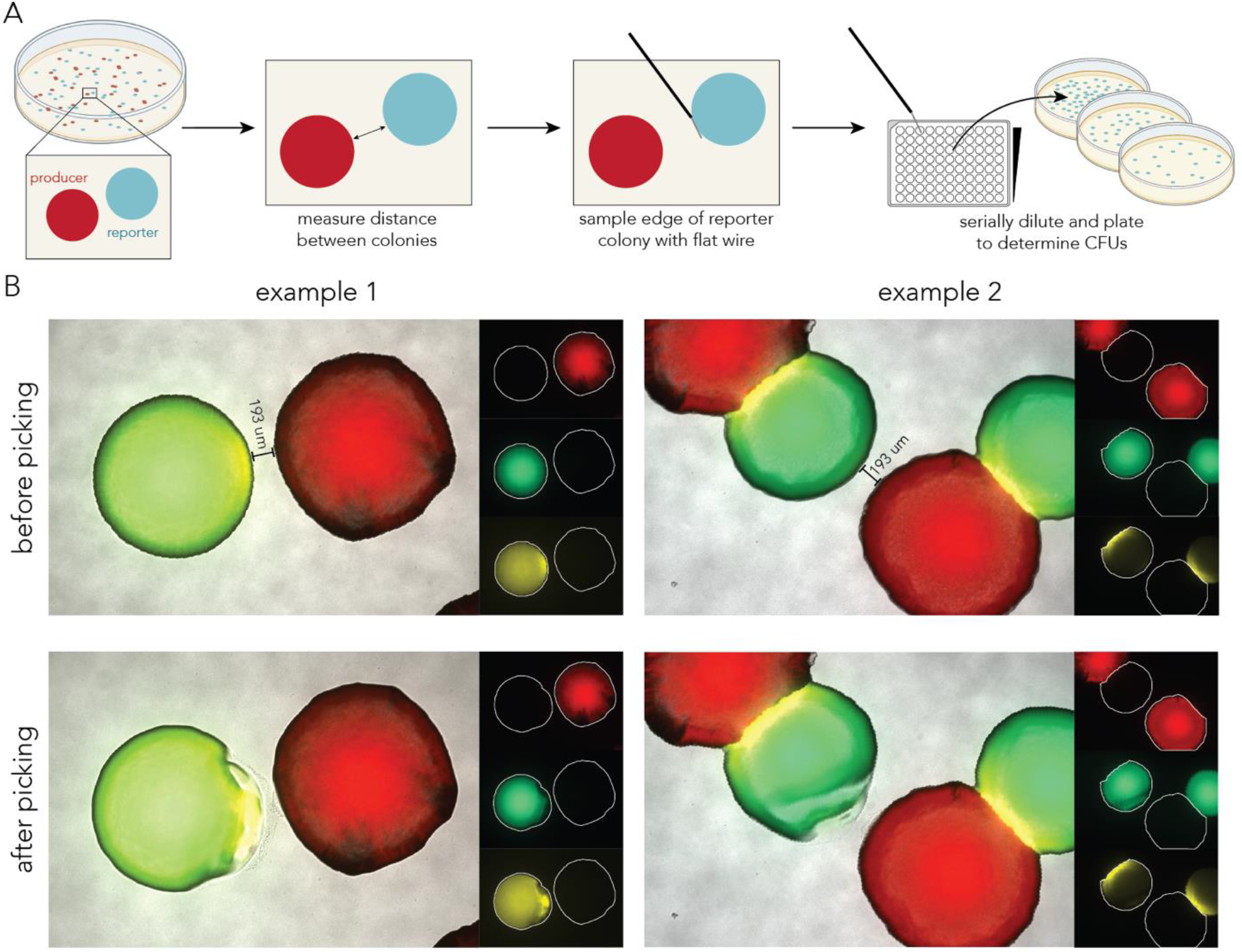
Validation of strain purity in the edge of non-contacting colonies. (A) Outline of the sampling and plating approach used to test for strain purity: non-contacting reporter and pks^+^ colony pairs were identified and the distance between them was measured. Then, the reporter colony was sampled with a fine needle and the sample was suspended in buffer for serial dilution and plating. After overnight growth, plates with highly dense yet separate colonies were inspected with a fluorescent microscope to evaluate strain purity (median colony number was 7,500). (B) Representative images of two non-contacting colony pairs before and after fine needle sampling of the reporter colony. After fine needle sampling of the reporter colony, we confirmed that the pks^+^ colony remained undisturbed by microscopy imaging (lower panels show only reporter colonies are disrupted while pks^+^ remained unchanged).

**Supplementary Table 1.** Strain purity in non-contacting colonies.

| pks+ strain | colony pair | distance between colonies (um) | # mCherry colonies | total colonies on plate | CFUs picked |
| --- | --- | --- | --- | --- | --- |
| engineered (BAC) | 1 | 211.0878559 | 0 | 6713 | 472000 |
| engineered (BAC) | 2 | 65.39144619 | 0 | 7617 | 948000 |
| engineered (BAC) | 3 | 196.5908064 | 0 | 7315 | 1898400 |
| engineered (BAC) | 4 | 218.0115838 | 0 | 6675 | 1424000 |
| engineered (BAC) | 5 | 141.7858862 | 0 | 10387 | 1284800 |
| engineered (BAC) | 6 | 250.630535 | 0 | 9619 | 1169600 |
| engineered (BAC) | 7 | 316.371115 | 0 | 10171 | 796000 |
| natural (EcN) | 1 | 109.1133998 | 0 | 7568 | 872000 |
| natural (EcN) | 2 | 61.83338299 | 0 | 7609 | 896000 |
| natural (EcN) | 3 | 207.0858625 | 0 | 4067 | 1300800 |
| natural (EcN) | 4 | 130.9372108 | 0 | 3662 | 668800 |
| natural (EcN) | 5 | 119.9792786 | 0 | 6802 | 672000 |

